# Utility of nasal swabs for assessing mucosal immune responses towards SARS-CoV-2

**DOI:** 10.1101/2023.07.12.548630

**Authors:** Ericka Kirkpatrick Roubidoux, Pamela H. Brigleb, Kasi Vegesana, Aisha Souquette, Kendall Whitt, Pamela Freiden, St. Jude Investigative Team, Amanda Green, Paul G. Thomas, Maureen A. McGargill, Joshua Wolf, Stacey Schultz-Cherry

## Abstract

SARS-CoV-2 has caused millions of infections worldwide since its emergence in 2019. Understanding how infection and vaccination induce mucosal immune responses and how they fluctuate over time is important, especially since they are key in preventing infection and reducing disease severity. We established a novel methodology for assessing SARS-CoV-2 cytokine and antibody responses at the nasal epithelium by using nasopharyngeal swabs collected longitudinally before and after either SARS-CoV-2 infection or vaccination. We then compared responses between mucosal and systemic compartments. We demonstrate that cytokine and antibody profiles differ markedly between compartments. Nasal cytokines show a wound healing phenotype while plasma cytokines are consistent with pro-inflammatory pathways. We found that nasal IgA and IgG have different kinetics after infection, with IgA peaking first. Although vaccination results in low nasal IgA, IgG induction persists for up to 180 days post-vaccination. This research highlights the importance of studying mucosal responses in addition to systemic responses to respiratory infections to understand the correlates of disease severity and immune memory. The methods described herein can be used to further mucosal vaccine development by giving us a better understanding of immunity at the nasal epithelium providing a simpler, alternative clinical practice to studying mucosal responses to infection.

**Teaser:** A nasopharyngeal swab can be used to study the intranasal immune response and yields much more information than a simple viral diagnosis.

## Introduction

In 2019, the SARS-2 coronavirus (SARS-CoV-2) pandemic began in Wuhan, China and quickly spread across the globe. The primary route of infection with SARS-CoV-2 is through the inhalation of respiratory droplets, with infections typically beginning at the mucosal surface of the nasal cavity^1,2^. The spike protein is responsible for viral entry via the angiotensin-converting enzyme 2 (ACE2) receptor on nasal epithelial cells^2^. It is also the target of antibody responses, with those targeting the receptor binding domain (RBD) being the most neutralizing^3, 4^. Shortly after infection, antibodies towards the RBD arise with IgA and IgG detectable around 9 days post-infection^5^.

Immunoassays have been developed as tools to study immune responses to infection and vaccination, however, they are focused on serological reactions to the virus. Serological assays are important and have aided in our understanding of anti-SARS-CoV-2 immunity and supported the development of SARS-CoV-2 vaccines. It is understood that the immune response can be compartmentalized, with mucosal responses differing from systemic^6, 7^. Additionally, mucosal immunity within the upper respiratory tract (URT) is a key factor in preventing and controlling infections^8–11^. Typically, saliva or nasal washes are collected as representative samples of the URT mucosal compartment. These types of samples do capture secretions from mucosal surfaces; however, they come with some caveats. Nasal washes are highly invasive, leading to participant hesitancy. Saliva is a non-invasive sample type, but saliva is not representative of the nasal mucosa, and its proximity to the gingiva can lead to a more intermediate phenotype between a mucosal and systemic sample^12^. The large volumes collected may also dilute the signal, and most studies do not control sample-to-sample variability. Finally, no studies to date have examined the longitudinal kinetics of nasal responses toward SARS-CoV-2.

To fill this gap in knowledge and further our understanding of innate and humoral immunity towards SARS-CoV-2 in the nasal cavity, we designed a series of experiments to measure cytokines and antibodies in nasopharyngeal (nasal) swabs collected longitudinally from individuals enrolled in our St. Jude Tracking of Viral and Host Factors Associated with COVID-19 (SJTRC) cohort study^13^. Participants were swabbed weekly in an institutional surveillance program to screen for SARS-CoV-2 infections. Nasal swabs were available from the baseline (pre-infection), acute, early convalescent, late convalescent, post convalescent, and late post convalescent phases of COVID-19 disease. After the release of the BNT162b2 (Pfizer) mRNA vaccines in late 2020, participants were offered vaccinations, providing an opportunity to measure nasal antibodies after vaccination. To determine the differences between nasal and systemic responses, cytokine and antibody levels were quantitated over time from the same participants. Our studies uncovered that longitudinal kinetics vary depending on infection or vaccination, antibody isotype, and viral antigen. We also noted that the mucosal and systemic responses are compartmentalized and have distinct profiles that persist long after infection or vaccination. Importantly, our work highlights that nasal swabs are a powerful, underutilized tool for further understanding nasal mucosal immunity.

## Results

### Study design and sample collection

Nasal swabs were collected weekly from participants enrolled in the SJTRC cohort study for asymptomatic monitoring and after infection to document clearance. We selected 48 individuals with RT-PCR confirmed SARS-CoV-2 infection (CT value < 40) and 26 vaccinated individuals (BNT162b2 [Pfizer] mRNA vaccination, 2 doses, three weeks apart) for these studies. Cohort characteristics are described in **Table 1**. Most infections were caused by SARS-CoV-2 B.1 lineage viruses and no severe illness was reported. Nasal swabs and a plasma sample were collected pre-exposure (baseline), and during the acute (1-21 days post infection, dpi), early convalescent (22-59 dpi), late convalescent (60-89 dpi), post convalescent (90-180 dpi), and late post convalescent (>180 dpi) phases of infection (**Fig. 1**). Additionally, nasal swabs and plasma were available from vaccinated individuals prior to vaccination and then during the 22-56 days post vaccination (dpv), 57-89 dpv, 90-180 dpv, and >180 dpv periods. None of the included vaccinated individuals were diagnosed with SARS-CoV-2 during this period. This study design provided us with an opportunity to investigate mucosal cytokine and antibody responses longitudinally compared to pre exposure levels. The collection of baseline nasal swabs and plasma is unique to this study and enabled us to make better inferences with the data, since baseline immune responses can vary amongst people.

**Table 1.** Cohort characteristics. The age range, sex distribution, race, and ethnicity of individuals in the infected (n=48) and vaccinated (n=26) cohorts. IQR stands for inter-quartile range with the first and third quartiles listed.

| Characteristic |  | Infection<br>(n=48) | Vaccinated<br>(n=26) |
| --- | --- | --- | --- |
| Age (Median [IQR]) |  | 43.00 [34.25, 53.00] | 41.00 [34.00, 55.75] |
| Sex (%) | Female | 40 (83.3) | 18 (69.2) |
|  | Male | 8 (16.7) | 8 (30.8) |
| Race (%) | Asian | 1 (2.1) | 4 (15.4) |
|  | Black/African American | 12 (25.0) | 3 (11.5) |
|  | WhiteCaucasian | 35 (72.9) | 19 (73.1) |
| Ethnicity (%) | Hispanic | 1 (2.1) | 0 (0.0) |
|  | Non-Hispanic | 40 (83.3) | 23 (88.5) |
|  | Other, Non-Hispanic | 7 (14.6) | 3 (11.5) |

**Fig. 1.**
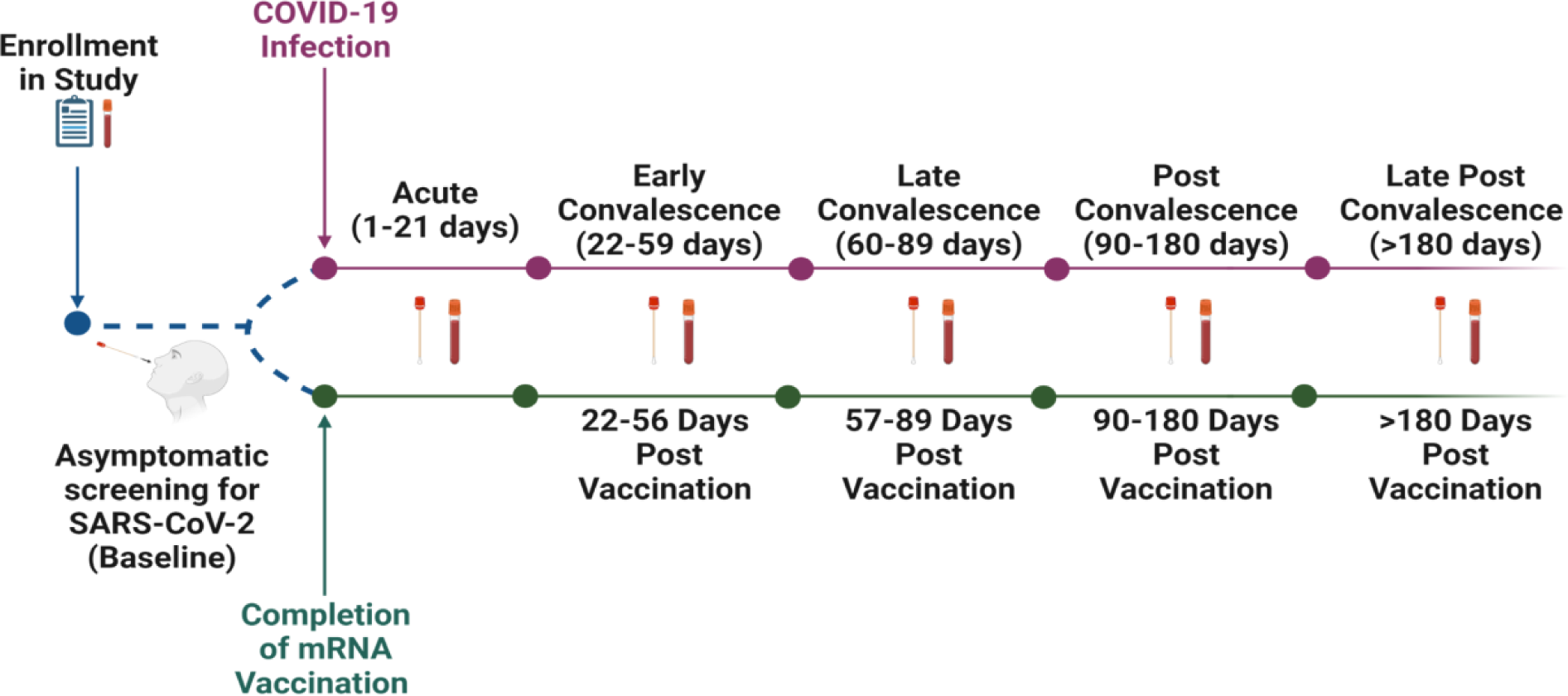
Study timeline. Individuals enrolled in the SJTRC study in early 2020. Upon enrollment, a blood sample and demographic information were collected followed by collection of weekly nasopharyngeal swabs as a part of a SARS-CoV-2 employee asymptomatic screening program. If someone tested positive prior to becoming vaccinated, they were included in the infected cohort. After testing positive, nasal swabs and plasma were collected during the acute, early convalescent, late convalescent, post convalescent, and late post convalescent phases of infection. If individuals managed to remain SARS-CoV-2 negative before receiving two doses of the Pfizer mRNA BNT162b2 vaccine, they were included in the vaccination cohort. These individuals also provided nasal swabs and plasma at 22-56 days post vaccination (dpv), 57-89 dpv, 90-180 dpv, and >180 dpv.

Nasal swab quality was assessed using RNase P qPCR. All nasal swabs with a Ct value below 40 were included in the study. Nasal swabs were then handled as depicted in **Figure 2**. To reduce swab-to-swab variability, total protein concentration was determined, and nasal swab material was diluted to a protein concentration of 0.5 mg/mL for subsequent assays. These steps ensured that each nasal swab was standardized for downstream experiments and allowed for more direct comparisons between samples. We then used these nasal swabs to assess longitudinal nasal cytokine and antibody kinetics in our infected and vaccinated cohorts.

**Fig. 2.**
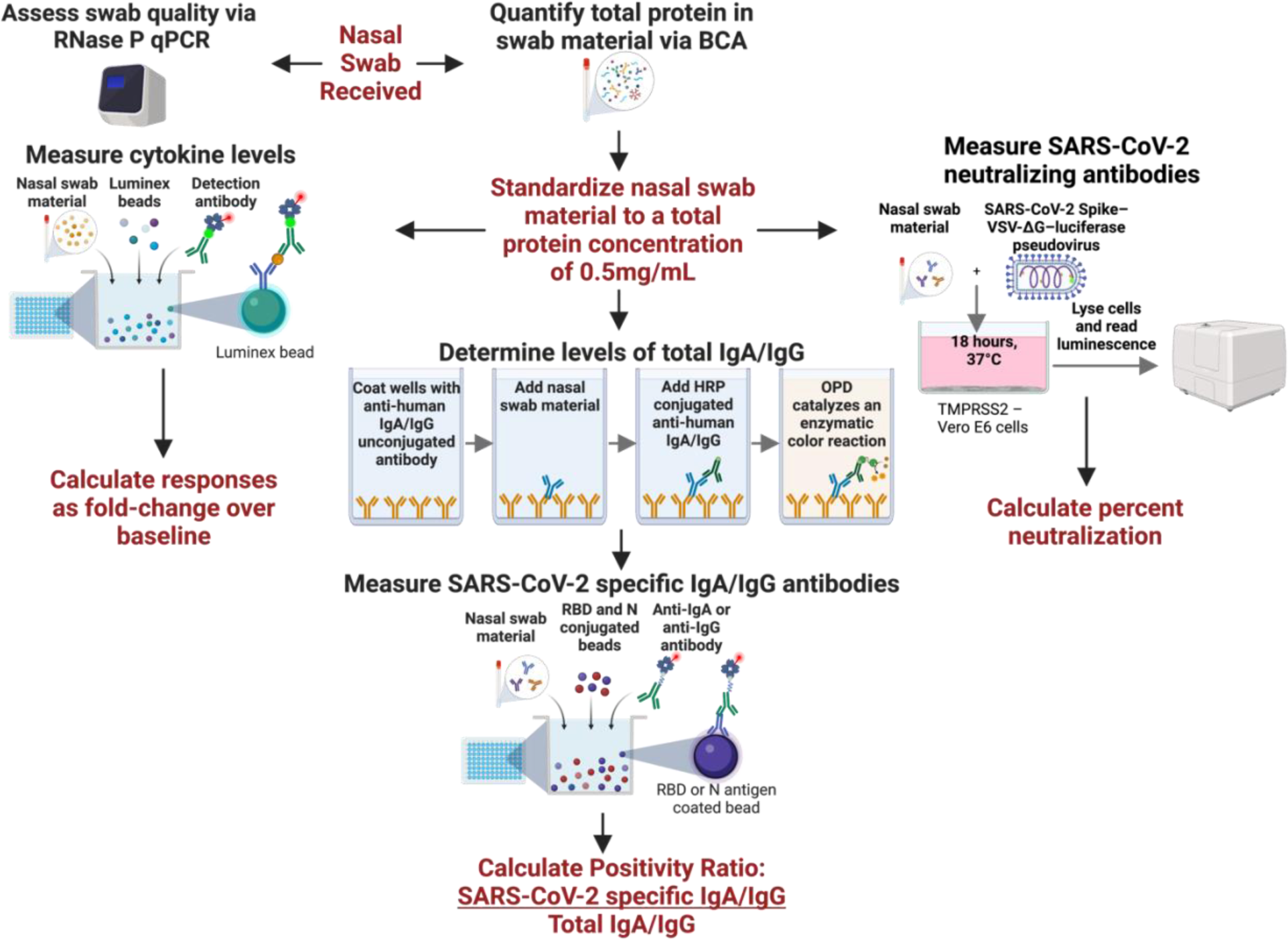
Methodology for measuring innate and adaptive mucosal immune responses from a single nasal swab. Upon receipt, nasal swabs were thawed and aliquoted. One aliquot was used to assess swab quality by the presence of RNase P. A second aliquot was used to determine total protein concentration using BCA. Nasal swabs were diluted to a standardized concentration of 0.5mg/mL for downstream assays to account for the differences in total protein. Cytokines were measured using a Luminex kit with streptavidin-PE conjugated detection antibody. We reported cytokine values as a fold-change over baseline. We determined total IgA and IgG levels using an ELISA with anti-human IgA or IgG as a capture antibody. A second, HRP-conjugated, anti-human IgA or IgG was used to detect IgA or IgG captured from nasal swab samples. The total peak area under the curve was calculated and used as the variable for total IgA or IgG levels. SARS-CoV-2 specific antibodies were measured using a Luminex based kit with streptavidin-PE conjugated anti-human IgA or IgG secondary antibodies. A “positivity ratio” was calculated by dividing antigen specific IgA/IgG by total IgA/IgG. Neutralizing antibodies were determined using a SARS-CoV-2 Spike-VSV-ΔG-luciferase pseudovirus. Nasal swab material was incubated with the virus for 1 hour prior to infecting confluent TMPRSS2 cells. The following day, cells were lysed and luminescencewas measured. Percent neutralization was calculated for each swab by comparing the nasal swab + virus luminescence to virus only luminescence.

### Cytokine and chemokine responses are distinct between nasal and systemic compartments

Cytokine levels are commonly assessed in the blood to determine systemic levels of inflammation. Prior studies, including one from this cohort^14^, have shown elevated systemic levels of specific cytokines, including IL-1Ra and IL-8, have been previously associated with an increased risk of severe disease and poor outcomes in persons with SARS-CoV-2 infection^15–18^. However, little is known about the mucosal immune response to SARS-CoV-2 infection and whether the mucosal immune responses correlate with systemic immune responses, especially at later time points. These are important to understand the long term impact of an upper-lower respiratory infection on mucosal immunity that may impact the susceptibility and severity to other respiratory infections. In this study, both mucosal (nasal) and systemic (plasma) immunity to natural SARS-CoV-2 infection were assessed by multiplex Luminex analysis to determine the levels of 31 different cytokines/chemokines. Since baseline levels differed (**Supplementary Fig. 1**), each acute or convalescent time point was normalized to their baseline.

Several plasma cytokines had an increased fold change following infection with the most robust increases observed with CXCl10, TNFα, IL-10, and IL-1RA, corroborating previously published data^19–22^. To investigate whether mucosal immune responses would trend similarly to systemic responses, we used a similar Luminex Cytokine Human Panel on nasal swab samples diluted to 0.5 mg/ml. Although we detected a smaller percentage of cytokines in the nasal swabs compared to the plasma, 12/30 compared to 30/30 in the plasma (**Fig. 3A & 3B**), there was an opposite trend to that observed in the plasma. Instead of the overall increase in cytokines seen in the plasma, we observed a decrease in several cytokine levels at the acute time point in the nasal swabs, including CCL2, IL1-RA, and IL-8, which were elevated in the plasma in the acute stage. This suggests that inflammatory immune responses are more concentrated systemically as the infection has migrated to other sites of infection (e.g., lungs or gastrointestinal tract) at those time points. Ingenuity pathway analysis of the cytokines up or down-regulated in the nasal swabs were consistent with a wound healing phenotype while those in the plasma with pathogen-induced inflammatory signaling pathways (**Fig. 3A & 3B**).

**Fig. 3.**
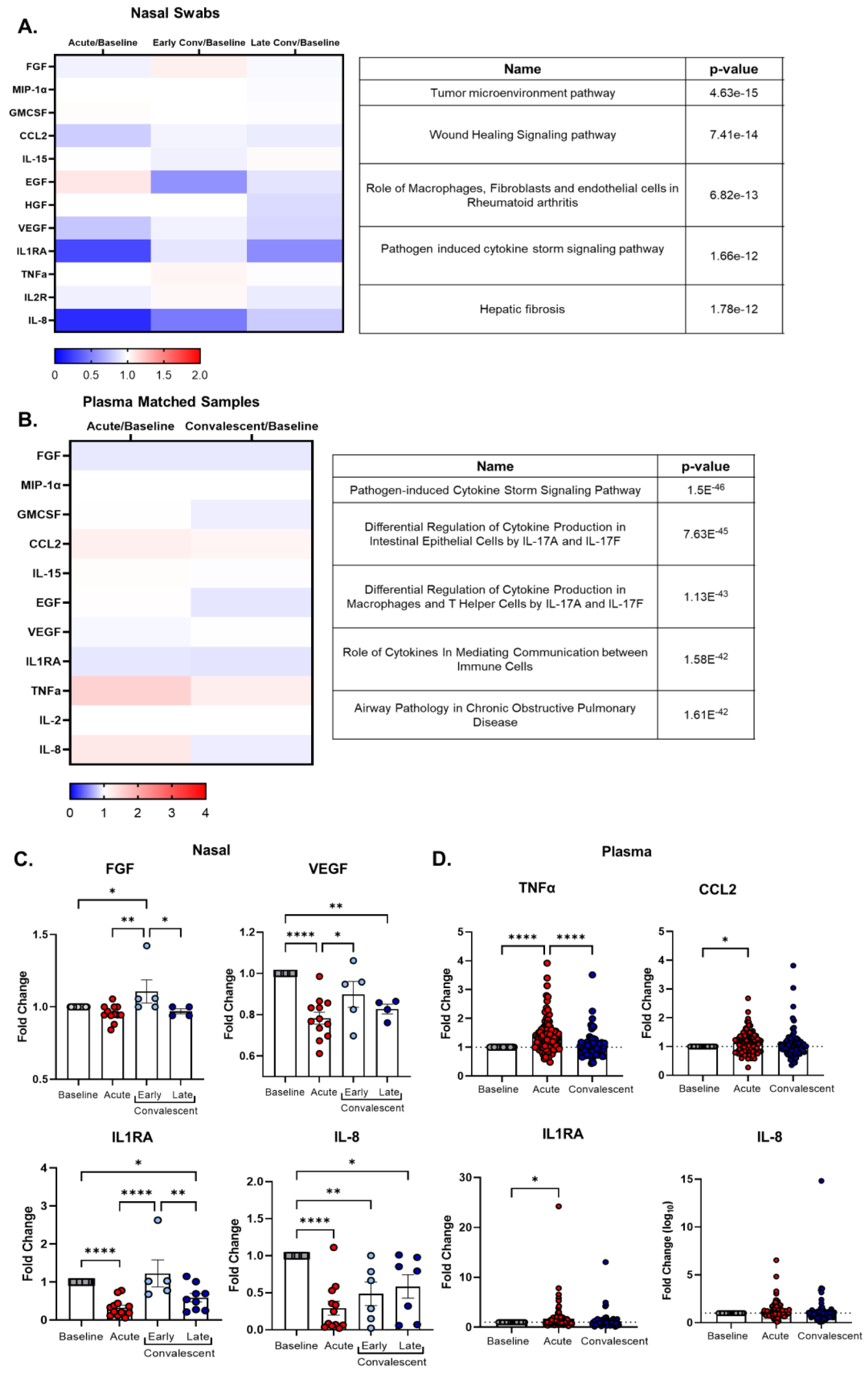
SARS-CoV-2 infection alters cytokine responses differentially in the plasma and nasal cavity over time. Nasal swabs or plasma samples were collected at various times-post testing positive for SARS-CoV-2 and a baseline sample for nasal and plasma pre-infection was used for normalization. Cytokines were assessed by multiplex Luminex assay. **(A, B)** Heat map of the median cytokine fold changes response to each person’s baseline value to account for human variation for nasal swabs **(A)** or plasma samples **(B).** Convalescent stage was split into early (days 21-62) and late (>62 days) post-infection to study the longitudinal impact of SARS-CoV-2 infection on mucosal cytokine responses **(A).** Ingenuity pathway analyses using predetermined signaling pathways on cytokines that were up or downregulated were assessed for both the nasal and plasma **(A, B). (C)** Fold change from baseline in acute, early, or late convalescent for cytokines FGF, VEGF, IL1RA, and IL-8 from nasal swabs. **(D)** Fold change from baseline in acute or convalescent from plasma for TNFα, CCL2, IL1RA, and IL-8. Heat maps and subsequent statistical analyses were conducted in GraphPad Prism version 9. Statistical analyses include a One-way ANOVA with Tukey’s Multiple Comparisons test (**D)**. * p < .05, ** p < .01, *** p < .001, **** p < .0001. Plasma, n=96 for baseline, acute and convalescent; nasal swabs, n=28 (baseline), n=12 (acute), and n=21 (total early + late convalescent). Note, not all individuals had cytokine levels detected in the nasal cavity at baseline or post-infection, which were excluded from this analysis.

One strength of these studies is the availability of longitudinal samples, allowing us to assess the impact of infection and vaccination on long-term systemic and mucosal immune responses. While no significant differences were observed in the plasma (**Fig. 3D**), we found that there were still changes in cytokine expression in the nasal cavity weeks following infection. In our early convalescent timepoint, there were several cytokines that were upregulated including FGF and IL2R (**Fig. 3B and 3D**). In late convalescence, we found that most cytokines were still downregulated in the nasal cavity, including IL-1RA, IL-8, and VEGF (**Fig. 3B and 3D**). These data suggest that infection with SARS-CoV-2 may alter cytokine responses, specifically at the mucosal surface, long term, which may have implications in reinfection and susceptibility to other respiratory infections.

### Development of high-throughput methodology for characterizing longitudinal mucosal antibody responses using nasal swab

After observing the stark differences between cytokine and chemokine expression in nasal and systemic compartments, we next wanted to evaluate whether they translated to differences in antibody expression. Also, it is important to understand how nasal antibody levels rise and fall after SARS-CoV-2 exposure or vaccination to determine the longevity of memory immune responses. To do this, we mapped the longitudinal nasal responses of both infected and vaccinated SJTRC participants as depicted in **Fig. 2**. Total IgA and IgG present in each nasal swab was measured using an enzyme-linked immunosorbent assay (ELISA). Area under the curve (AUC) analyses were performed and used as the value for total IgA or IgG. SARS-CoV-2 RBD and N-specific antibodies were detected using a multiplex Luminex assay, and the average mean fluorescent intensity (MFI) for each sample was reported. Baseline nasal swabs were used to establish background signal. To normalize the levels of SARS-CoV-2 specific antibodies in proportion to total IgA or IgG present in a sample, we calculated a positivity ratio of antigen-specific IgA or IgG to total IgA or IgG.

Importantly, we were able to confidently detect SARS-CoV-2 specific IgA and IgG at the nasal epithelium using nasal swabs. Within the infected cohort, anti-RBD and -N IgA titers peak early after infection and then steadily decline (**Fig. 4**). Anti-RBD and -N IgG titers rise and remain at moderate levels for an extended period. In most cases, anti-RBD and N antibodies return to baseline levels by late post-convalescence. The exception is anti-RBD IgG titers, which increase between the post convalescence and late post convalescence phases for several individuals. These individuals had received their first dose of the BNT162b2 mRNA vaccine in the intervening period, which explains the discrepancy. Of note, they are not included in the vaccinated cohort.

**Fig. 4.**
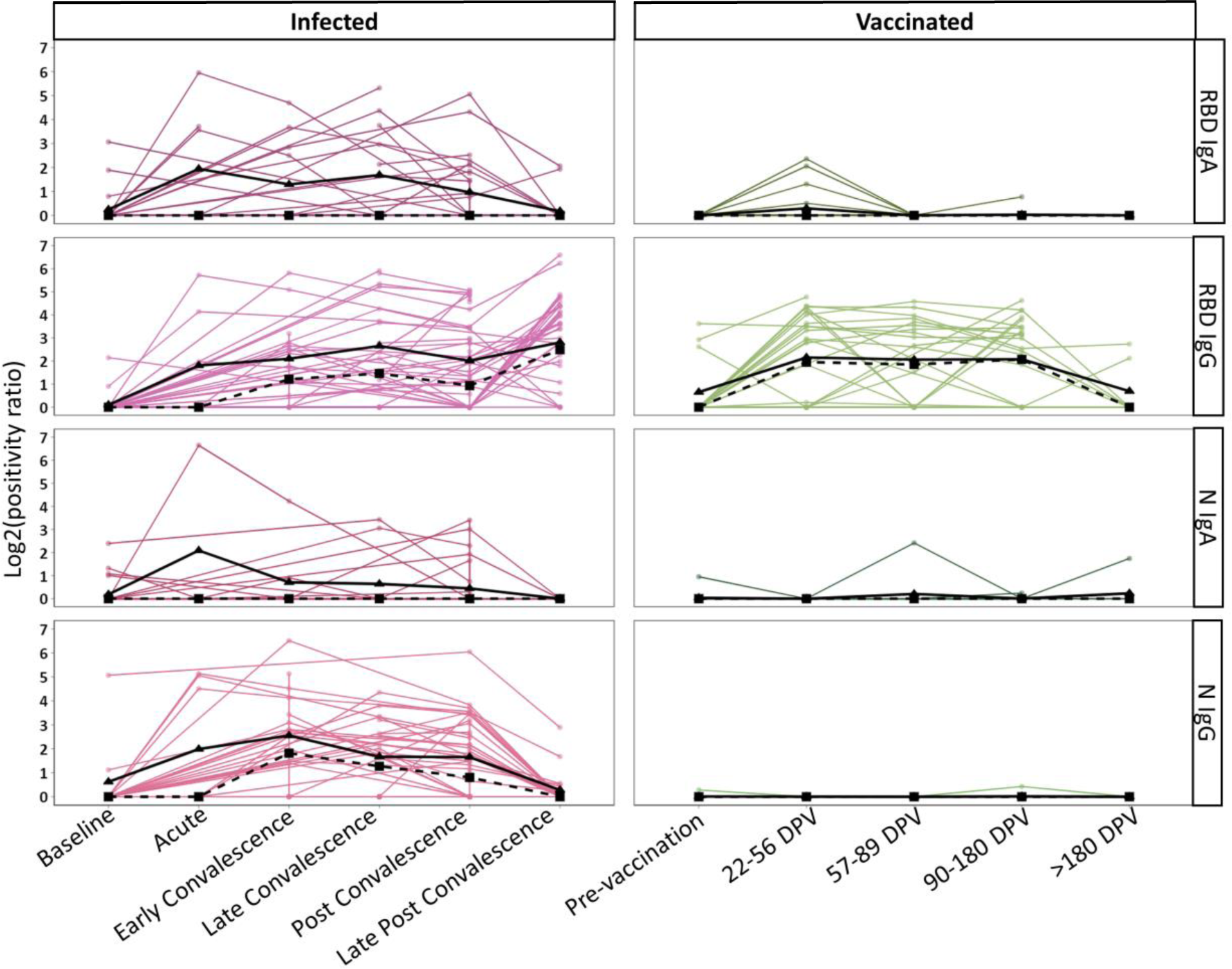
Longitudinal kinetics of mucosal anti-SARS-CoV-2 IgA and IgG in infected and vaccinated individuals. The responses of infected (left) and vaccinated individuals (right) are shown. Positivity ratios are shown for anti-RBD IgA, anti-RBD IgG, anti-N IgA, and anti-N IgG. Collection timepoints are listed for each cohort as either a phase of infection or days post vaccination (DPV). The solid black line on each graph represents the mean response and the dotted line represents the median response at each time point.

Within the vaccinated cohort, we observed a small peak of anti-RBD IgA at 22-56 dpv (**Fig. 4**). On average, this peak was half of the response observed for the corresponding time post infection (early convalescence). Their levels return to baseline by 57-89 dpv. Anti-RBD IgG titers rose to similar levels as infected individuals and remained stable for up to 180 dpv. As expected, anti-N titers were negligible. Infection leads to an IgA and IgG response to both SARS-CoV-2 antigens evaluated while vaccination appears to only induce a strong anti-RBD IgG response.

Using cohort data previously collected about the plasma response to SARS-CoV-2^23–25^, we mapped the matched plasma antibody levels of participants **(Supplemental Fig. 2)** and observed that IgA antibody kinetics between the compartments are different, with IgA peaking during early convalescence in plasma. Additionally, IgA responses seem to last longer in plasma from both infected and vaccinated participants. IgG responses were more uniform between the nasal and systemic compartments. This may be due to the mechanisms of IgA and IgG induction and transport since IgA is produced locally in the nasal cavity while IgG is bi-directionally transported between the two compartments^26–28^. This data highlight that the location and timing of sample collection could greatly impact the observed antibody response. Nasal responses are more readily detected after exposure, while systemic responses persist longer.

### Assessing the nasal swab neutralization activity

Serological data show that neutralizing antibodies are a correlate for protection from SARS-CoV-2 infection and severe disease^3, 4^. Studies using other mucosal samples such as saliva and nasal washes have shown that antibodies in the URT can be neutralizing^11, 29, 30^. It is important to be able to detect and measure neutralization activity of antibodies at the nasal epithelium, especially since this is the primary site of infection. We chose nasal swabs with the top 10% anti-RBD IgA and IgG positivity ratios to assess whether neutralizing antibodies are detectable at the nasal epithelium using a SARS-CoV-2 spike VSV-ΔG-luciferase pseudovirus. We observed that nasal swabs from infected individuals had a higher average neutralizing activity compared to the vaccinated cohort (**Fig. 5**). Within the infected cohort there was a wider range of neutralizing capacity, with few nasal swabs completely neutralizing the virus. Only one nasal swab within the vaccinated cohort had elevated levels of neutralizing antibodies. This data suggests that while the quantity of anti-RBD IgG antibodies is similar between infected and vaccinated cohorts, the quality is not. Upon further analysis, we noted that neutralizing nasal swabs had higher anti-RBD IgG positivity ratios compared to anti-RBD IgA positivity ratios, suggesting that neutralization is more IgG driven **(Supplemental Fig. 3 and Supplemental Table 1)**. This is a consideration to make when evaluating new, mucosal vaccine responses as it will be important for them to induce long-lived IgG responses that are effectively trafficked to the nasal epithelium.

**Fig. 5.**
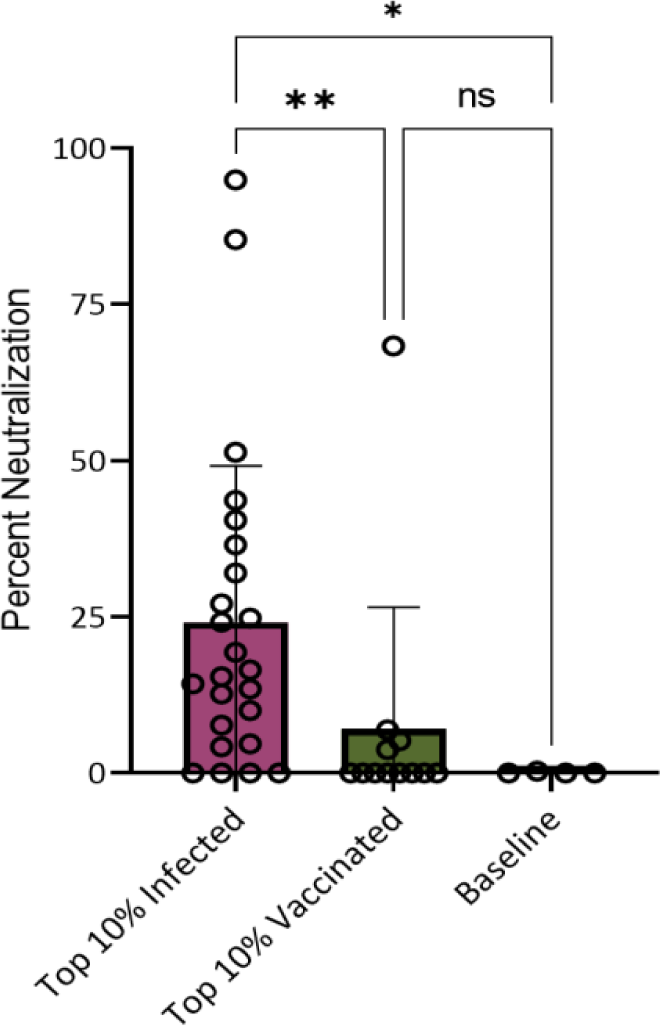
Neutralization activity is higher in nasal swabs from infected individuals. Nasal swabs with the top 10% anti-RBD IgA and IgG positivity ratios were selected for neutralization assays. Percent neutralization of the swab material at a concentration of 0.5mg/mL total protein is shown. For the infected cohort N=24, for the vaccinated cohort N=12, and a final N of 4 of baseline samples was included. Each sample was run in duplicate. One baseline sample was removed prior to statistical analyses after being identified as an outlier through a ROUT test in PRISM 9. Kruskal-Wallis multiple comparisons were used to detect significant differences between groups. *Indicates P=0.0299, **indicates P=0.0057, and ns stands for non-significant.

### Longitudinal IgG responses do not exhibit compartmental bias

Other studies, which used a variety of mucosal samples, have reported compartmental bias between mucosal and systemic compartments^6, 7, 29, 31, 32^. However, the comparison of longitudinal responses between nasal and systemic components has not yet been made. Within this study, data of antibody levels in the nasal cavity and plasma were collected differently (positivity ratio of antigen specific antibodies to total antibody levels vs ELISA determined optical density (OD) of antigen specific antibodies, respectively), making it difficult to directly compare responses between the compartments. We used a ranking system to compare whether an individual had a similar overall antibody response between both compartments. To do this, we calculated AUCs of the total nasal and plasma responses across all time points for each person within the study. We then ranked each positive individual from lowest to highest response and compared whether those with a high nasal rank also have a high plasma rank. Individuals with no response were given a rank of 0. We then graphed this data in a scatterplot and divided it into 4 quadrants: the top left represents those who had the highest 25% plasma responses, top right represents those who had the highest 25% of both plasma and nasal responses, bottom right represents those who had the highest 25% nasal responses, and the bottom left represents those who had the lowest 25% of both plasma and nasal responses (**Fig. 6**).

**Fig. 6.**
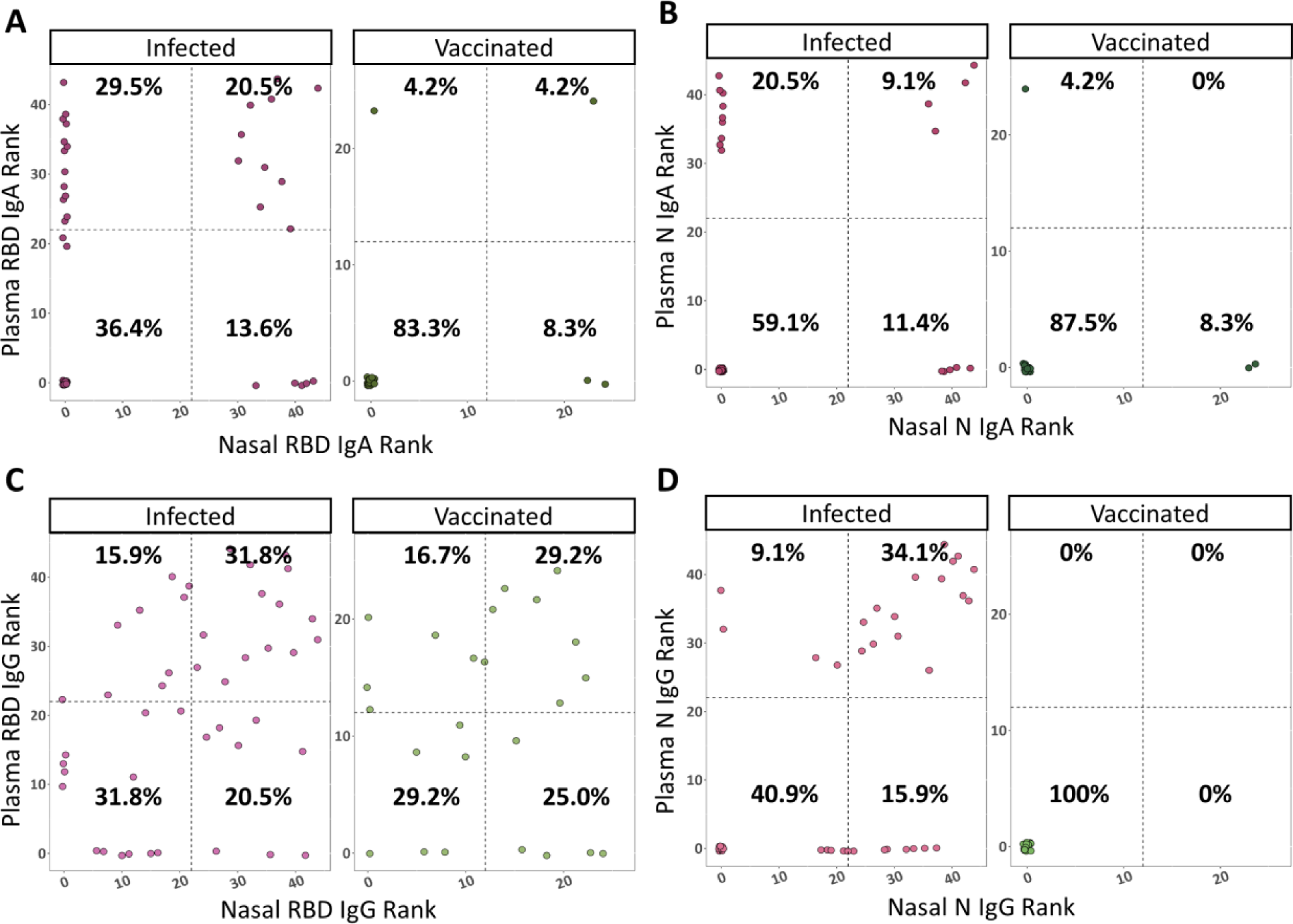
Anti-SARS-CoV-2 responses have compartmental bias. Each person’s longitudinal nasal and plasma response was summarized using AUC analyses, calculated using R software. AUCs were ranked from lowest to highest, with the highest rank indicating the best response. Individuals with no response were given a rank of 0. These ranks are presented in scatterplots, with the dotted lines dividing them into 4 quadrants representing high plasma responses (top right), high nasal and plasma responses (top left), high nasal responses (bottom right), and low responders (bottom left). The percentage of people within each quadrant is listed on the graphs. **(A)** Plasma vs nasal anti-RBD IgA ranks. **(B)** Plasma vs nasal anti-N IgA ranks. **(C)** Plasma vs nasal anti-RBD IgG ranks. **(D)** Plasma vs nasal anti-N IgG ranks.

In all cases, most individuals fell into the lower left quadrant for both nasal and plasma responses. This may be because participants did not report any severe disease, which is known to induce a stronger antibody response^33, 34^. A larger percentage of infected individuals ranked higher for plasma IgA than nasal IgA, regardless of antigen (**Fig. 6A-B**). This observation is driven by the fact that IgA persists longer in plasma, increasing the longitudinal AUC. Little IgA was observed in the vaccinated cohort; therefore, it is difficult to conclude whether the response was biased towards plasma or nasal responses. We observed better overall responses towards RBD compared to N, especially within IgA.

IgG responses had little compartmental bias, with the highest percentage of people ranking in the top 25% of plasma and nasal responses (**Figure 6C-D**). Anti-RBD IgG responses ranked very similarly between infected and vaccinated cohorts, which is corroborated by their comparable antibody kinetics. This data suggests that to get a complete picture of anti-SARS-CoV-2 IgA responses, both nasal and plasma compartments are important. However, overall responses to IgG are quite similar regardless of compartment or route of antigen exposure.

## Discussion

This study demonstrates that nasal swabs can be used for more than diagnostic testing. By collecting baseline samples and standardizing nasal swab material, we established methodology that elucidated the longitudinal nasal anti-SARS-CoV-2 cytokine and antibody response. The data shown here highlights the importance of studying immunity at the infection site and systemically. The same pro-inflammatory cytokines upregulated systemically and were significantly downregulated at the nasal epithelium. The differences between nasal and systemic antibodies were less stark, however we did see that induction kinetics and longevity of IgA differ between compartments. The nasal response seems more short-lived compared to a plasma response. This is important to consider because the time of sample collection can play a role in influencing the levels of a respective antigen-isotype combination. Correlates for protection between nasal and systemic compartments will be vastly different depending on the timing and type of sample taken.

IgA seems to be more compartmentalized compared to IgG. The reasons behind this observation are both biologically and experimentally driven. First, IgA is produced in mucosal tissues and a secretory version is transported to mucosal surfaces through pIg receptors, meaning that IgA is inherently more mucosal than IgG^28, 35, 36^. Additionally, IgA is more short-lived compared to IgG so there is a higher likelihood of IgG being detected at later time points regardless of compartment^28, 35, 36^. Other reasons for this observation can be due to experimental nuances. IgA quaternary structure is diverse and can be monomeric, dimeric, multimeric, or can contain a secretory signal (sIgA). The anti-human IgA secondaries used for this study may be biased towards plasma and not sIgA, making responses appear more compartmentalized than they really are. Interestingly, the extent of compartmental bias is different depending on longitudinal collection time highlighting the importance of long-term human cohort studies when investigating mucosal and systemic immunity.

We found that IgG kinetics were similar between infected and vaccinated cohorts regardless of bodily compartment. However, neutralizing IgG was only present in the infected cohort. This is important to note because it indicates that neutralizing antibodies induced by vaccination are not being trafficked to the nasal epithelium. Our finding that neutralizing mucosal antibodies correlates with nasal IgG is unique. Other studies suggest that mucosal IgA is critical for neutralizing mucosal responses^5, 37, 38^. However, these studies used nasal washes or saliva as mucosal samples, and therefore are not measuring nasal epithelium. Our data suggests that IgA at the nasal epithelium is not playing as strong of a role in neutralization as IgA present in mucosal fluids. It underscores the importance of sample type when examining neutralizing mucosal responses. However, it should be noted that a luciferase-based pseudovirus platform is not as sensitive as some other SARS-CoV-2 neutralization assays and the use of live virus may increase the likelihood of identifying neutralizing IgA containing nasal swabs^39^.

We established the first study to examine longitudinal nasal antibody kinetics and confirmed existing reports of distinct cytokine responses. Additionally, longitudinal sampling revealed that SARS-CoV-2 infection shifts the cytokine profile at the nasal epithelium and can take months to return to baseline levels, which may impact reinfection with SARS-CoV-2 or other respiratory pathogens. Finally, we uncovered important new information about nasal antibody kinetics that show compartmental bias can be observed at the antibody level as well. Kinetics differ between antibody isotype and site of collection, indicating that long term studies will provide the best information in terms of understanding the antibody response to SARS-CoV-2 as it is a very dynamic process. Infected individuals in our cohort reported mild to no clinical disease, which was reflected in their low antibody titers both mucosal and systemically. This additionally made it difficult to draw significant conclusions as to how the reported cytokine data influenced antibody outcomes. If studies using nasal swabs are continued in cohorts with stronger antibody responses, we may be able to use nasal swabs to detect biomarkers for poor disease outcomes or correlates of protection. Future work involving mucosal immunity should also consider investigating immunity within the nasal cavity as it gives a better picture of what is occurring at the site of infection compared to other mucosal samples. Additionally, nasal swabs are more ideal mucosal samples as they are sampling the correct anatomical location while being only mildly invasive.

## Methods

### Study design and sample handling

Participants in SJTRC provided written consent to participate in the institutional review-board approved, prospective study^13^. This study began with the collection of a blood sample (baseline), and the completion of a demographic survey, summarized in **Table 1**. As part of St. Jude COVID-19 employee surveillance, participants were swabbed weekly until staff vaccination became ubiquitous, and swabs collected prior to diagnosis or vaccination were selected as baseline samples. Longitudinal nasal swabs and blood were collected after PCR-confirmed infection or after the second dose (“completion”) of the Pfizer mRNA BNT162b2 vaccine (**Fig. 1**). While nasal swabs and plasma were collected during the same time periods, they were not necessarily collected concurrently. Additionally, not all nasal swab samples have a matched plasma sample. Nasal swabs were collected using FLOQ Swabs (COPAN, Cat No. 520CS01) and placed in 1mL of Viral Transport Media (DMEM with 0.25% FBS (Fetal Bovine Serum)) at 4°C and were stored at - 80°C after collection. SARS-CoV-2 diagnostics were performed by the clinical microbiology laboratory at St. Jude the day of nasal swab collection. The remaining sample was kept at -80°C until received by our laboratory. Samples were thawed, RNA was immediately isolated, and remnants stored at 4°C for antibody and cytokine assays. Plasma was isolated from whole blood and stored at -80°C until needed for antibody and cytokine assays and kept at 4°C after thawed. Data are managed using an electronic database hosted at St. Jude (REDCap). The infected cohort consisted of 48 individuals and the vaccinated cohort consisted of 26 individuals.

### RNase P qPCR

Human nasal swab samples were inactivated with 350 µls RLT buffer (Qiagen, Cat No. 79216) containing 1% β-mercaptoethanol for a minimum of 10 minutes. Following manufacturer’s recommended directions, RNA was extracted using a RNeasy Mini kit (Qiagen, Cat No. 74106) and assessed on a Nanodrop 2000. Four microliters of RNA were added to a 16ul master mix containing nuclease-free water (Teknova, Cat No. W3330), TaqMan Fast Virus 1-Step Master Mix (ThermoFisher, Cat No. 5555532) and a commercially prepared RNAse P primer/probe combination (IDT, Cat Nos. 10006827, 10006828, 10007061, 10006829, 10011568) to quality test for the human RNAse P (RPP30) gene. A portion of the RPP30 gene was used as a positive control (IDT, Cat No. 10006626) and nuclease-free water was used as a negative control. A qRT-PCR assay was run on a BioRad CFX96 Real Time System with cycling conditions: 25°C for 2 mins, 50°C for 15 mins, 95°C for 3 mins, followed by 45 rounds of 95°C for 15 secs, 55°C for 30 secs, data acquired. Ct values under 38 were considered positive. RNA extractions and qPCRs were performed in singlet. Only one nasal swab was removed from the study due to a high RNase P Ct value.

### Measuring total protein

The concentration of total protein in each nasal swab was determined using the Pierce™ BCA Protein Assay Kit (ThermoFisher, Cat No. 23225) according to the manufacturer’s microplate procedure. Briefly, neat nasal swab material and a 1:5 dilution of material in phosphate buffered saline (PBS) were added to a clear, 96-well plate. Optical density (OD) at 562nm was read using BioTek Synergy2 plate reader and Gen5 (v3.09) software. All samples and standards were performed in duplicate, with averages used for calculations. Standard curve calculations were done in excel and concentrations were determined based on the average of both the neat and 1:5 dilution of nasal swab material, unless one of these values was out of the range used to determine the standard curve (>2mg/mL or <0.025mg/mL). Once total protein concentrations were calculated, all nasal swabs were diluted to a standard concentration of 0.5mg/mL in sterile 1xPBS and kept at 4°C for all downstream experiments.

### Cytokine and chemokine assays

Cytokine levels were measured from plasma or nasal swab samples that included a baseline measurement per individual. Cohort plasma acute cytokine data were also used for a previous study^14^. Nasal swabs were pre-diluted to 0.5 mg/ml for consistency with other protein analyses in this study. Cytokines were measured using the Human Cytokine Magnetic 30-Plex Panel (Invitrogen, Cat No. LHC6003M) and plates were read using a Luminex200 machine with xPONENT software (v4.3). Each sample was run in duplicate, and the average read was used for subsequent analyses. Sample exclusion from analyses included failure of detection for all cytokines and having no baseline value for comparison. Ingenuity pathway analysis (IPA) was used to identify pathways cytokines that were up or downregulated during the acute phase relative to baseline.

### Total IgA and IgG ELISAs

Total IgA and IgG ELISAs were performed using 384-well flat-bottom MaxiSorp plates (ThermoFisher, Cat No. 464718) coated with either an unconjugated anti-human IgA (Novusbio, Cat No. NB7441) or an unconjugated anti-human IgG (Novusbio, Cat No NBP1-51523) antibody at 2μg/μL in 1xPBS (**Fig. 2**). Once coated, plates were left overnight at 4°C. Plates were washed 4 times with PBS containing 0.1% Tween-20 (PBS-T) using the AquaMax 4000 plate washer system. After washing, plates were blocked with PBS-T containing 0.5% Omniblok non-fat milk powder (AmericanBio, Cat No. AB10109-01000) and 3% goat serum (Gibco, Cat No. 16210-072) for 1 hour at room temperature. The wash buffer was removed, and plates were tapped dry. Nasal swab material at 0.5mg/mL was serially diluted 1:3 in blocking solution and run in duplicate. Recombinant human IgA (abcam, Cat No. ab91025) or recombinant human IgG (abcam, Cat No. ab91102) was also diluted to 5μg/mL and ran in duplicate on each plate for quality control. D After 2 hours at room temperature, plates were washed 4 times with PBS-T. Anti-human IgA HRP (Novusbio, Cat No NBP1-73613) diluted 1:2000 or anti-human IgG HRP (Creative Biolabs, Cat No. MOB-0361MC) diluted 1:5000 was then added to the plates and left to incubate for 1 hour at room temperature. Plates were washed 4 times with PBS-T and developed using SIGMAFAST™ OPD (Sigma-Aldrich, Cat No. P9187) for 10 minutes at room temperature. The developing reagent was inactivated using 3M hydrochloric acid (Fisher Scientific, Cat No. A144-212). Plates were read at 490nm using a BioTek Synergy2 plate reader and Gen5 (v3.09) software. For each plate, an upper 99% confidence interval (CI) of blank wells OD values was determined and used as the Y= value in an area under the curve (AUC) analysis in PRISM 9. AUC was determined for each nasal swab and used as the denominator in positivity ratio calculations.

### IgA specific nasal antibodies

SARS-CoV-2 anti-RBD and -N IgA antibody levels were determined using 2 kits due to a shortage of supplies while conducting experiments. To prevent kit-to-kit variability, samples were run concurrently on both kits and MFIs (Mean Fluorescent Intensity) were correlated. Additionally, positive control antibodies for RBD (InvivoGen, Cat No. srbd-mab6) and N (GenScript, Cat No. A02090) were included at high (30µg/mL) and low (0.01µg/mL) concentrations to each plate to monitor for plate-to-plate variability. We observed a strong correlation between MFI values for samples and control antibodies between the kits (r 0.9951, p <0.0001) **(Supplemental Fig. 4)**. The first kit used was the Milliplex® SARS-CoV-2 antigen panel 1 IgA assay (Millipore Sigma, Cat No. HC19SERA1-85K) with the Wuhan-1 strain RBD and N proteins included. The second kit used was the Bio-Plex Pro Serology Reagent Kit (Bio-Rad, Cat No. 12014777), with human IgA positive and negative controls (Bio-Rad, Cat No. 12014775), Bio-Plex SARS-CoV-2 Wuhan-1 strain RBD and N coupled beads (Bio-Rad, Cat Nos. 12015406 and 12014773), and Bio-Plex Pro Human IgA detection antibody (Bio-Rad, Cat No. 12014669). For both kits, the manufacturers’ instructions were followed, except that nasal swab material was diluted to 0.5mg/mL in PBS. The protocol for the Milliplex® SARS-CoV-2 kit involved incubating nasal swab material with RBD- and N conjugated beads in the dark for 2 hours at room temperature, shaking. The beads were then washed three times using a handheld magnetic separation block (EMD Millipore, Cat No. 40-285). Next, PE-anti-human IgA conjugate was added to each well and incubated in the dark for 90 minutes at room temperature, shaking. Beads were washed again three times and then resuspended in sheath fluid and stored at 4°C overnight, shielded from light. The protocol for the Bio-Plex kit involved incubating RBD- and N conjugated beads with nasal swab material in the dark for 30 minutes at room temperature, shaking. Plates were washed 3 times using a handheld magnetic separation block and then human IgA detection antibody was added and incubated in the dark for 30 minutes, shaking. Next, plates were washed 3 times and SA-PE was added for 10 minutes, in the dark and shaking. Finally, beads were washed again three times and then resuspended in sheath fluid and stored at 4°C overnight, shielded from light. The following day, plates were read using a Luminex200 machine with xPONENT software (v4.3) and data was analyzed as “qualitative” using kit specific recommended plate layout and settings. All samples were measured in duplicate.

### IgG specific nasal antibodies

SARS-CoV-2 anti-RBD and N IgG antibody levels were determined using the Milliplex® SARS-CoV-2 antigen panel 1 IgG assay (Millipore Sigma, Cat No. HC19SERG1-85K) with the Wuhan-1 strain RBD and N proteins included. The protocol was followed as described in the kit instructions, except that nasal swab material diluted to 0.5mg/mL in 1xPBS instead of assay buffer. Control antibodies for RBD (InvivoGen, Cat No. srbd-mab12) and N (AcroBiosystems, Cat No. NUN-S41) were added at high (30µg/mL) and low (0.01µg/mL) concentrations to each plate to monitor for plate-to-plate variability. Briefly, nasal swab material was incubated with RBD- and N conjugated beads in the dark for 2 hours at room temperature, shaking. The beads were then washed three times using a handheld magnetic separation block. Next, PE-anti-human IgG conjugate was added to each well and incubated in the dark for 90 minutes at room temperature, shaking. Beads were washed again three times and then resuspended in sheath fluid and stored at 4°C overnight, shielded from light. Plates were read using a Luminex200 machine with xPONENT software (v4.3) and data was analyzed as “qualitative” using kit recommended plate layout and settings. All samples were measured in duplicate.

### Calculation of positivity ratios

To account for non-specific signal from nasal swab material, we used baseline swabs to establish a positive/negative cutoff MFI for each antigen-isotype pair. All baseline MFI values for RBD IgA, RBD IgG, N IgA, and N IgG were individually averaged, and the top 99% confidence interval (3 standard deviations above the mean) was used as the antigen-isotype specific cutoff value. All MFIs below the cutoff value were given a negative value to ensure that only positive samples would have high positivity ratios. Next, the AUC of total IgA or total IgG for each nasal swab (determined via ELISA) was used as the denominator to calculate the positivity ratio (MFI/AUC). A positivity ratio of 1 or lower was considered negative.

### Determining plasma IgA and IgG antibodies

Cohort plasma IgA and IgG antibodies were determined using an ELISA and used for previous studies^23–25^. Briefly, plates were coated with 1.5µg/mL of Wuhan-1 RBD or 1µg/mL Wuhan-1 nucleoprotein (produced in-house) and left overnight at 4°C. Next, plates were blocked with 3% milk in PBS-T for 1 hour at room temperature. Plates were washed three times and a 1:50 dilution of plasma in 1% milk was added to the plate for 1.5 hours at room temperature. Next anti-human IgA HRP (Novusbio, Cat No. NBP1-73613) diluted 1:2000 or anti-human IgG HRP (Creative Biolabs, Cat No. MOB-0361MC) diluted 1:10,000 was added for 30 minutes at room temperature. Plates were washed again and developed using SIGMAFAST™ OPD (Sigma-Aldrich, Cat No. P9187) for 8 minutes at room temperature and then stopped using 3M hydrochloric acid (Fisher Scientific, Cat No. A144-212). Plates were read at 490nm using a BioTek Synergy2 plate reader and Gen5 (v3.09) software. OD was reported and anything 2-fold above negative control plasma (OD 0.15) was considered positive. Plasma samples were tested in duplicate.

### Neutralization Assays

Neutralization assays were performed using a SARS-CoV-2 spike VSV-ΔG-luciferase pseudovirus that was generated as previously described^39^. Approximately 24 hours prior to the assay, VeroE6/TMPRSS2 cells (XenoTech, Cat No. JCRB1819) were plated at 2.5×10^4^ cells per well in a clear, 96-well tissue culture treated plate in DMEM (Corning, Cat No. 10-013-CV) supplemented with 5% FBS (Sigma, Cat No. F2442) (D-5). The following day, nasal swab material was diluted in D-5 media to 0.5mg/mL. The SARS-CoV-2 spike VSV-ΔG-luciferase pseudovirus was diluted to 250 infectious units (IU) and this was incubated with nasal swab material for 1 hour at 37°C in 5% CO_2_. The VeroE6/TMPRSS2 cells were then washed 1 time with 1xPBS and the virus+swab material mixture was immediately added. Plates were placed at 37°C in 5% CO_2_ and left for 16-18 hours (overnight). The following day, Luc-Screen™ Extended-Glow Luciferase buffers 1 and 2 (ThermoFisher, Cat No. T1035) used according to manufacturer’s instructions. Luminescence was measured using the BioTek Cytation3 plate reader with 1 sec integration time and analyzed with Gen5 (v3.09) software. Percent neutralization was calculated using the following equation: 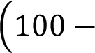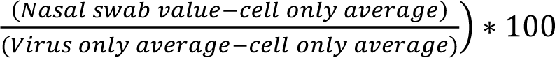 Virus only and cell only averages were calculated for each plate individually. As a control, a neutralizing monoclonal antibody towards SARS-CoV-2 (SinoBiologicals, Cat No. 40592-R0004) was included on each plate. All samples were run in duplicate. Percent neutralization, IgA positivity ratio, and IgG positivity ratio of each nasal swab tested are listed in **Supplemental Table 1**.

### Statistical analysis

Data was managed using the software REDCap and visualized using R or PRISM 9.0. For cytokine and chemokine analyses, heat maps and subsequent statistical analyses were conducted in GraphPad Prism version 9 as described in figure legends. Statistical analyses include a One-way ANOVA with Tukey’s Multiple Comparisons test. For the neutralization data, outliers were identified using the ROUT test and significant differences between groups were detected using a Kruskal-Wallis multiple comparisons (with standard parameters) test in PRISM 9.0 as described in figure legends.

## Acknowledgements

We would like to thank all SJTRC participants for their invaluable contribution to this study. Additionally, we would like to thank the staff of St. Jude Children’s Research Hospital and Lauren Rowland for their contributions.

