## Supplemental Materials for "Utility of nasal swabs for assessing mucosal immune responses towards SARS-CoV-2"

**Supplemental Table 1: Percent neutralization of nasal swabs.** Nasal swab percent neutralization at 0.5mg/mL, time point of collection, RBD IgA/IgG positivity ratios, and cohort are listed.

| Swab ID | Percent Neutralization at 0.5mg/mL | Timepoint | IgA RBD Positivity Ratio | IgG RBD Positivity Ratio | Cohort |
|---|---|---|---|---|---|
| 636 | 95 | Late Convalescent | 6.8 | 11.6 | Infected |
| 937 | 85 | Post-Convalescent | -0.2 | 95.7 | Infected |
| 839 | 68 | 90-180 days after completion | 0.7 | 7.8 | Vaccinated |
| 758 | 51 | Post-Convalescent | -4.1 | 28.1 | Infected |
| 693 | 44 | Late Convalescent | -1.5 | 36.3 | Infected |
| 690 | 40 | Late Convalescent | -4.9 | 39.5 | Infected |
| 764 | 37 | Post-Convalescent | -7.3 | 32.1 | Infected |
| 899 | 32 | Post-Convalescent | -0.2 | 32.8 | Infected |
| 906 | 30 | Post-Convalescent | -0.7 | 74.3 | Infected |
| 645 | 27 | Early Convalescent | 6.2 | 0.4 | Infected |
| 763 | 25 | Late Convalescent | 12.5 | 54.9 | Infected |
| 652 | 19 | Acute | 10.8 | 2.9 | Infected |

| | | | | | |
|---|---|---|---|---|---|
| 646 | 17 | Post-Convalescent | 18.9 | 1.1 | Infecte d |
| 685 | 15 | Late Convalescent | 38.8 | 59.3 | Infecte d |
| 761 | 14 | Post-Convalescent | -4.8 | 119.1 | Infecte d |
| 691 | 13 | Post-Convalescent | -5.1 | 28.1 | Infecte d |
| 734 | 13 | Early Convalescent | 24.9 | 6 | Infecte d |
| 687 | 10 | Late Convalescent | 19.6 | -12 | Infecte d |
| 733 | 8 | Acute | 60.7 | -10.5 | Infecte d |
| 799 | 7 | 90-180 days after completion | 0 | 17.6 | Vaccin ated |
| 767 | 5 | 22-56 days after completion | 1.5 | 0 | Vaccin ated |
| 615 | 5 | Acute | 12.1 | -4.5 | Infecte d |
| 619 | 4 | Early Convalescent | -13.9 | 55.3 | Infecte d |
| 769 | 4 | 90-180 days after completion | -0.7 | 17.2 | Vaccin ated |
| 681 | 0 | Baseline | -0.7 | -0.1 | Baselin e |
| 672 | 0 | Post-Convalescent | 32.2 | -1.9 | Infecte d |
| 747 | 0 | Acute | -20.4 | 51.7 | Infecte d |

| | | | | | |
|---|---|---|---|---|---|
| 748 | 0 | Early Convalescent | -6.4 | 33 | Infected |
| 873 | 0 | Post-Convalescent | -0.3 | 30.3 | Infected |
| 779 | 0 | 22-56 days after completion | 4.1 | 6.8 | Vaccinated |
| 837 | 0 | 22-56 days after completion | 0.4 | -0.1 | Vaccinated |
| 798 | 0 | 57-89 days after completion | 0 | 22.9 | Vaccinated |
| 802 | 0 | 22-56 days after completion | -0.3 | 19.9 | Vaccinated |
| 807 | 0 | 22-56 days after completion | 0 | 19.3 | Vaccinated |
| 846 | 0 | 22-56 days after completion | 3.2 | 19.3 | Vaccinated |
| 847 | 0 | 57-89 days after completion | -1.5 | 19.1 | Vaccinated |
| 856 | 0 | 90-180 days after completion | -0.1 | 23.7 | Vaccinated |
| 677 | 0 | Baseline | -0.5 | 0 | Baseline |
| 718 | 0 | Baseline | -0.4 | 0 | Baseline |
| 736 | 0 | Baseline | -1.4 | 0 | Baseline |

4

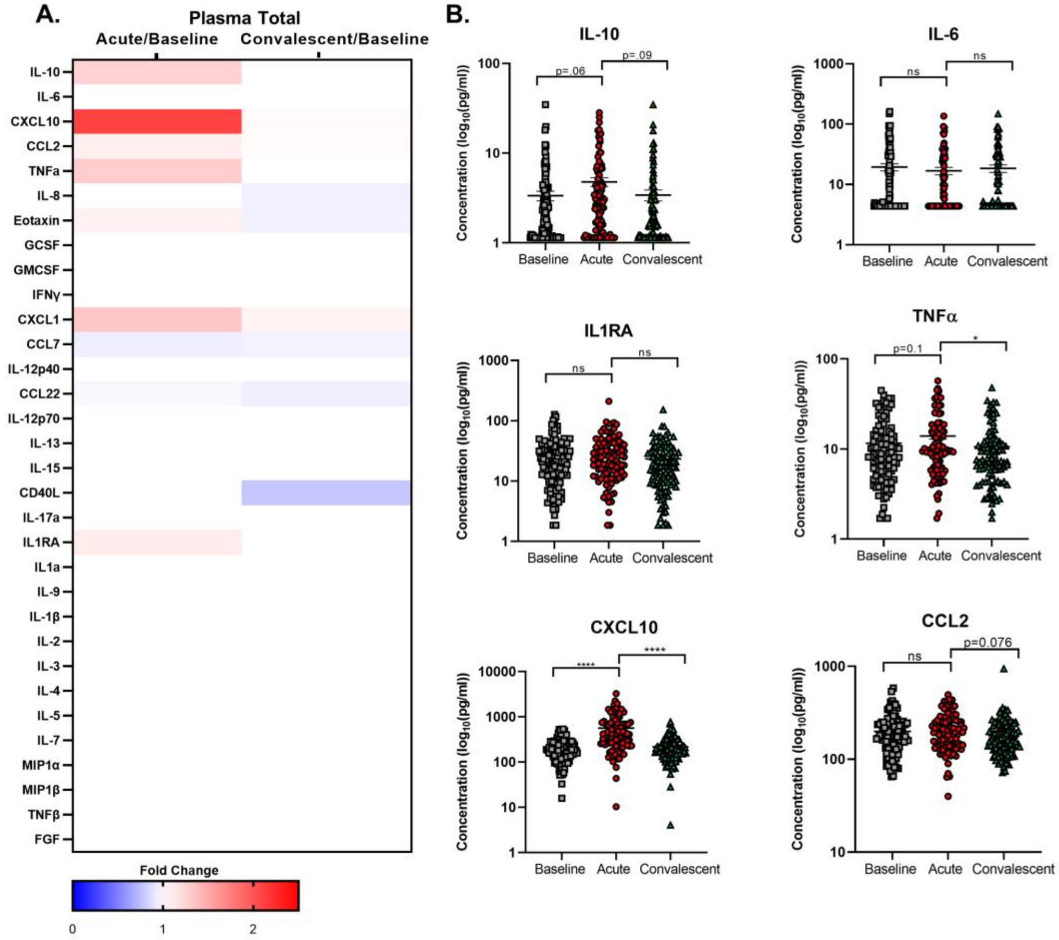


**Supplemental Figure 1. Systemic responses to SARS-CoV-2 natural infection.** Cytokine levels were assessed by multiplex Luminex at acute or convalescent time periods (N=95-96). **(A)** Total heat map of the median cytokine expression fold changes relative to baseline values. **(B)** Examples of plasma cytokine levels without normalizing to each person's own baseline value, demonstrating the high variability and range of values with limited trends or significance without normalization. Heat maps and subsequent statistical analyses were conducted in GraphPad Prism version 9. Statistical analyses include a One-way ANOVA with Tukey's Multiple Comparisons test **(B)** * $p < .05$, **** $p < .0001$.

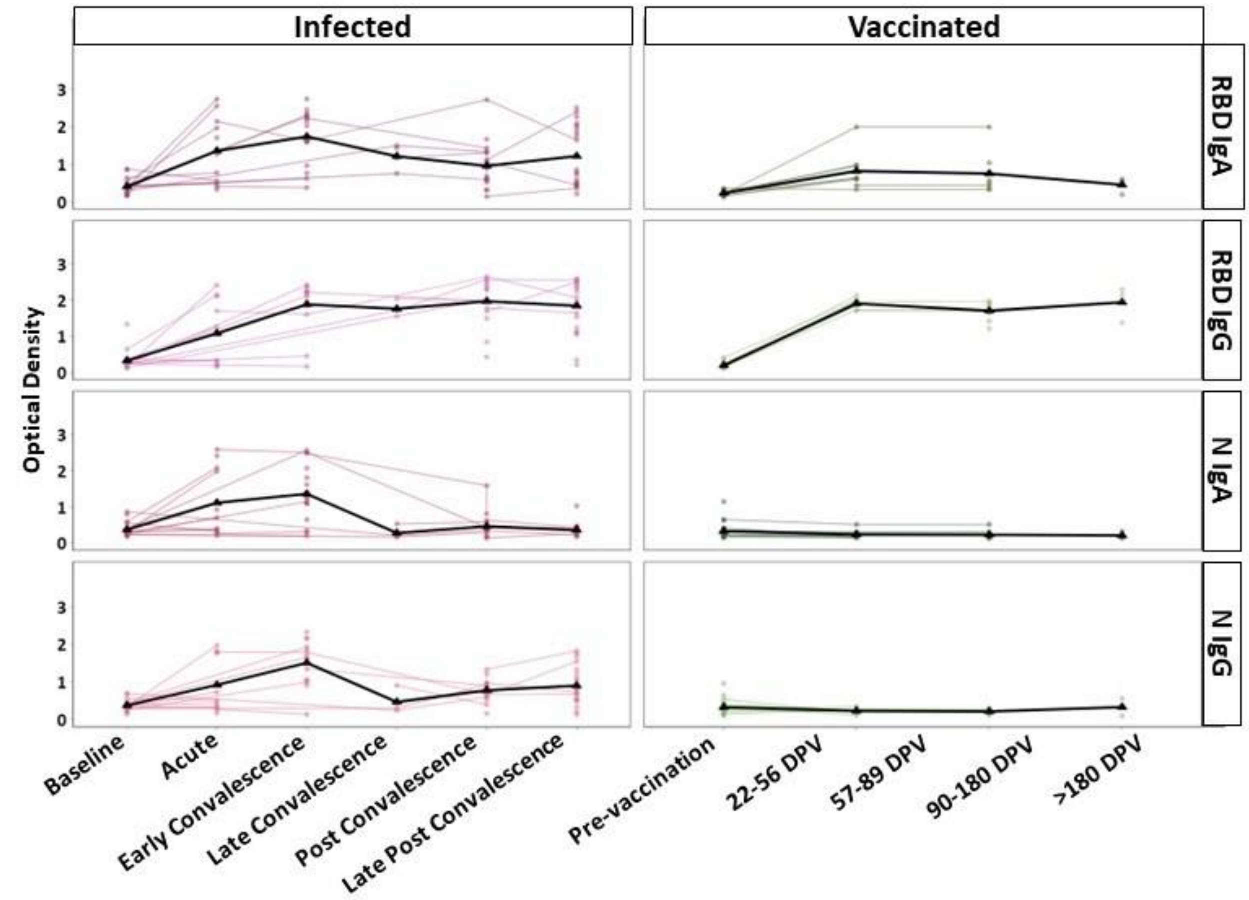


**Supplemental Fig. 2. Matched longitudinal plasma antibody kinetics.** The responses of infected (left) and vaccinated individuals (right) are shown. Positivity ratios are shown for anti-RBD IgA, anti-RBD IgG, anti-N IgA, and anti-N IgG. Collection timepoints are listed for each cohort as either a phase of infection or days post vaccination (DPV). The black line on each graph represents the mean response at each time point.

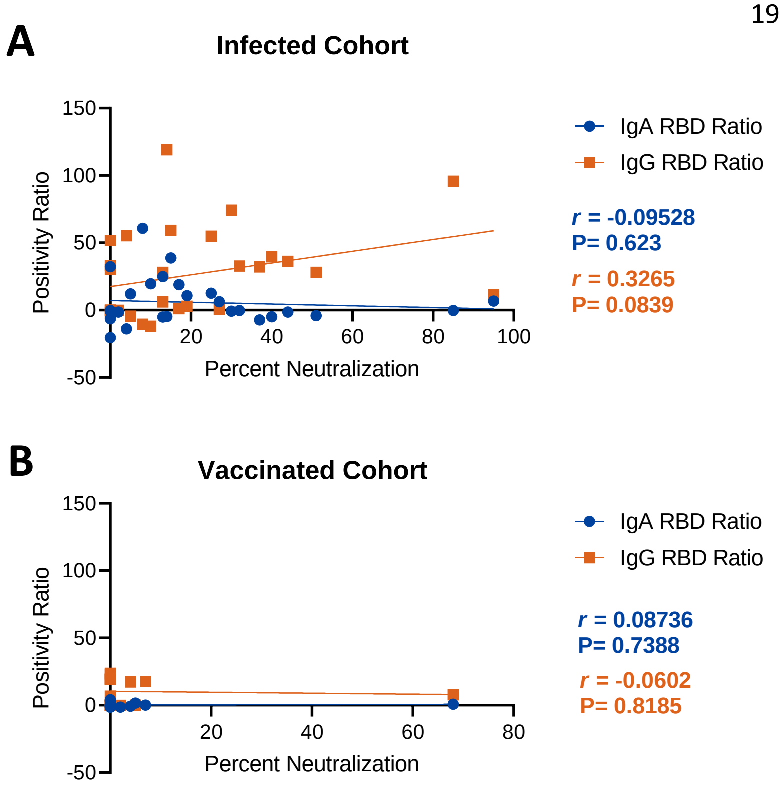


**Supplemental Fig. 3. Nasal swab neutralization is better correlated with anti-RBD IgG levels.** Scatterplots depict the correlation between percent neutralization and IgA or IgG positivity ratios for **(A)** infected and **(B)** vaccinated cohorts. IgA ratios are in blue while IgG ratios are in orange with the Pearson r and P value for each shown in the corresponding color. Additionally, simple linear regressions were performed, and the best-fit line is shown. Correlations were calculated using PRISM 9.0.

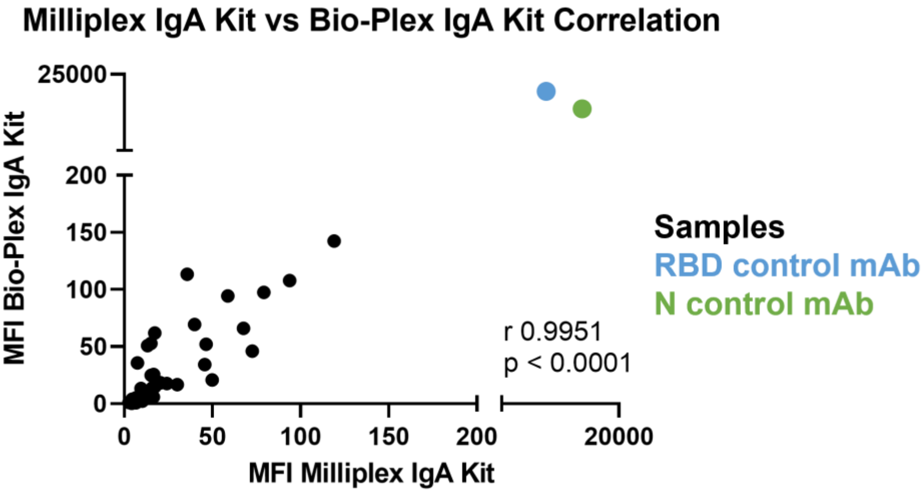

**Supplemental Fig. 3. Correlation between two different multiplex kits to measure antigen specific IgA in nasal swabs.** Kits were used following the manufacturer's instructions. Samples were diluted to 0.5mg/mL and MFI was measured using each kit. Monoclonal antibodies (mAbs) for RBD and N were also included. Pearson r correlation coefficient (0.9951) and p-value (<0.0001) are on the graph. Both were calculated using PRISM 9.0.
